# Hypothalamic dopamine neuron activity is modulated by caloric states and amphetamine abuse in zebrafish larvae

**DOI:** 10.1101/2024.01.31.578277

**Authors:** Pushkar Bansal, Erica E. Jung

**Affiliations:** Department of Mechanical and Industrial Engineering, The University of Illinois at Chicago, 842 W. Taylor St., Chicago, IL 60607, USA; Department of Bioengineering, The University of Illinois at Chicago, 851 S. Morgan St., Chicago, IL 60607, USA

**Keywords:** Food deprivation, Hypothalamic DA neurons, amphetamine, A11/A14 DA neurons, Zebrafish larva

## Abstract

Dopaminergic (DA) neuron activity is affected by reward and stress-inducing entities such as food and drugs of abuse, and different DA neuron populations can respond distinctly to these stimuli. Interaction between both stimuli significantly alters the dynamics of DA release in different DA populations. Additionally, these stimulating entities can affect the interconnections among different DA populations by impairing their correspondence with each other. However, limited studies have been performed that could point to the effect of interaction between AMPH and caloric states on DA neurons and their inter-correlation. This study explores the individual and interactive effect of two caloric states, ad-libitum fed (AL) and food deprived (FD), and acute exposure to a stimulant drug (amphetamine) in two different DA neurons in the hypothalamus of zebrafish larvae. We used a transgenic zebrafish line Tg(th2:GCaMP7s), which expresses a calcium indicator (GCaMP7s) in A11(Posterior Tuberculum) and a part of A14 (Caudal Hypothalamus and Intermediate Hypothalamus) DA populations located in the hypothalamus of the larval zebrafish. The larvae were subjected to acute FD and ad AL feeding followed by acute treatment with 0.7uM and 1.5uM doses of AMPH. We recorded calcium activity and quantified fluorescence change, activity duration, peak rise/fall time, and latency in the spikes of the DA neurons. Our results show that baseline DA neuron activity amplitude, spike duration, and correlation between inter- and intra-DA neurons were higher in the FD than in the AL state. Dose-dependent AMPH treatment further increased the activity intensity of the aforementioned parameters in the neuron spikes in the FD state. The DA activity correlation and spike latency were dose-dependently impaired in both DA populations. These results suggest that different DA populations in the brain exhibit a similar activity trend in response to caloric states and AMPH, where the AMPH-mediated intensity change in the activity was dose-dependent.

## Introduction

Dopaminergic neurons constitute a major part of the reward system in the brain and are located largely in the midbrain and forebrain nuclei. In mammals, several different populations of dopaminergic (DA) neurons (A8-A16) have been identified(Björklund & Dunnett, 2007). These populations were studied in response to various stimuli, mostly correlated with their specific function. DA neurons are primarily affected by reward and stress-inducing entities such as feeding/caloric states (food and hunger). Previous studies show that substantia nigra (SN) in the brain consisting of A9-type DA neurons showed an altered burst firing and spike frequency in response to food deprivation(Branch et al., 2013; GRENHOFF et al., 1988). In the A10 (Ventral Tegmental Area (VTA) and Nucleus accumbens (NAc)) and A12 (arcuate DA neurons), fasting increased dopamine neuron excitability in rodents(Roseberry, 2015; X. Zhang & Van Den Pol, 2016). Repercussions of hunger on A8 (retrorubral field (RRF)) type neurons were not directly studied, however, A8 neurons contribute to the mesolimbic reward circuitry consisting of VTA-A10 neurons(Gasbarri et al., 1996; Serafini et al., 2020). The DA neurons located in zona incerta (ZI) (A13) exhibited a significant decrease in activity in the food-deprivation state post-food retrieval in a recent study(Ye et al., 2023).

Addictive substances such as stimulants with abuse potential can also largely influence and increase DA activity in the brain (Calipari & Ferris, 2013). Amphetamine (AMPH) is one of the most widely used and abused substances (Kramer et al., n.d.). The effect of AMPH on DA activity was studied in the midbrain VTA-NAc-DA (A10) population (Vezina, 1988). In rats, the effect of AMPH on SN-A9 neurons showed that AMPH damaged the DA synaptic connection in SN and decreased striatal DA levels (Koirala et al., 2014). On the contrary, AMPH injection in SN-A9 neurons increased DA release in the pedunculopontine nucleus (Fougère et al., 2019; Ryczko et al., 2016). PeVN-A11 neurons lack DA transporters (DAT), and AMPH significantly increased DA activity in DAT lacking DA neurons (Giros et al., 1996; Koblinger et al., 2014). In PeVN-A14 neurons, DA neuron activity has not been studied directly. However, both A14 neuron activation and AMPH are known to increase locomotion, and chemogenetic inhibition of A14 neurons reduced AMPH-induced hyperlocomotion(Korchynska et al., 2022). This study suggests that AMPH increased A14 neuron expression. Taking these previous studies together, activities in RRF-A8 neurons, PeVN-(A11, A14, A15), and OB-A16 (Olfactory Bulb) DA neurons have not been studied in response to hunger (German & Manaye, 1993; Sánchez-González et al., 2005; Tillet et al., 1990; Vogt Weisenhorn et al., 2016). Additionally, activities in RRF-A8, PeVN-(A11, A15), ZI-A13, and OB-A16 DA neurons due to AMPH treatment remain unexplored. Moreover, besides midbrain VTA-NAc-A10 NAc, none of the populations were studied under the interactive effect of caloric states and AMPH (Stuber et al., 2002). Studying this interaction on DA neurons is important because caloric states, especially hunger, are known to potentiate amphetamine consumption, which could affect DA release.

DA populations in the brain are highly conserved across different animal species, including zebrafish (Barrios et al., 2020; Wasel & Freeman, 2020). Here, we used larval zebrafish, a vertebral teleost, to investigate the dopaminergic responses to food deprivation and a stimulant drug (AMPH). Zebrafish is a vertebral animal model gaining momentum in neuroscience research, especially due to their highly conserved brain structure and genome compared to mammals. Its small size (∼3mm) makes it easy to house for raising and handling during experimentation. Zebrafish larvae are optically transparent, which is advantageous in imaging and recording the activities inside the larval body, such as in the heart and brain (Bansal et al., 2023; Keller & Ahrens, 2015). We leveraged all these useful features, including optical transparency in the larvae, and recorded the change in DA neuron activity at a single neuron level using calcium imaging *in vivo* in real-time. To accomplish this, we used 6dpf larvae of a transgenic zebrafish line Tg(th2:GCaMP7s) expressing fluorescent activity in A11 (PT) and A14 (cH, iH) DA neurons in its hypothalamus. We treated the larvae with two different doses of AMPH (0.7uM and 1.5uM) while they were in ad libitum fed (AL) and food-deprived (FD) state. Our results show that DA neuron activity and correlation between both DA neuron populations were significantly increased in the FD state alone and with post-AMPH treatment in the FD state than in the AL state. On the other hand, latency between the neuron firing was significantly lower in the AL state than in the FD state after AMPH administration. Overall, our data, in accordance with the previous studies, indicate the potentiating effect of both hunger and its combination with AMPH on DA neuron activity across hypothalamic DA populations.

## Materials and Methods

### Zebrafish Maintenance

Parent transgenic zebrafish line Tg(th2:GCaMP7s) was outsourced and raised in our fish facility at the University of Illinois at Chicago. All adult fish tanks were stacked in the fish racking system (Aquaneering, Inc. San Diego, CA). The experiments conducted here were approved by the Association for Assessment and Accreditation of Laboratory Animal Care (AAALAC). All zebrafish larvae used here were 6dpf and were kept in the fish facility till they turned 6dpf. The fish facility was maintained at 28o C with a 14/10h light/dark cycle. Larvae were checked for any visible anomalies and were excluded from the study. The fish rack that constituted the fish water was maintained with a conductivity range of 600-800 us and a pH range of 7.2-7.8.

### Experimental Groups

The larvae were divided into two feeding groups: Ad Libitum (AL, n=20) and Food-deprived (FD, n=18). Ad Libtum larvae had unrestricted access to food since they turned 5dpf until the experimentation. Food-deprived larvae were never fed at any stage of raising and experimentation. Both feeding groups were treated with two different doses of amphetamine (AL 0.7uM, AL 1.5uM, and FD 0.7uM, FD 1.5uM). Initial dopamine neuron recording was performed for five continuous minutes in fish water (without amphetamine). After five minutes, the recording was stopped, and fish water was pipetted out and replaced with amphetamine solution. The neuron recording was again performed for another five minutes.

### Sample Preparation and Recording Material

For recording DA neuron activity, the individual larva was first paralyzed using 30 uL of 300uM stock pancuronium bromide (YC10036-1, 10MG, Millipore Sigma, WI, USA) to prevent any unwanted muscle twitching in the larvae to avoid error in recordings. After paralysis, larvae were completely immobilized in a 1.5% agarose gel solution drop and were covered with 0.5 ml fish water for 5 min baseline recording. After Baseline recording, the fish water was pipetted out and replaced with 0.5 ml d-amphetamine hemisulfate solution (Sigma Aldrich, MO). The drug was allowed to seep through the agarose drop for ∼5 mins, and treatment recording was commenced for 5 mins again. We used a 40x water immersion objective to visualize and record the neuron (40x/0.80 W, Olympus Corporation, Japan) mounted on an upright epifluorescence microscope setup consisting of a fluorescence excitation and illumination setup (X-cite 120 series, Excelitas Technologies Corp., Canada). The microscope was equipped with a GFP filter with 488nm excitation/ 507nm emission (Chroma Technology Corp, VT)). The setup was connected to an image visualization software(HC Image Live, Hamamatsu Photonics, Japan). The recording was made at 10ms exposure time with 35 frames per second. Recordings were opened in ImageJ, and a complete neuron cluster (3approx. n= 3-7 visible neurons/larva/DA population) (Fig. 1a) was selected from all three populations separately in the same frame using a polygon ROI tool along the edges of the cluster to avoid surrounding noisy regions.

**Figure 1.**
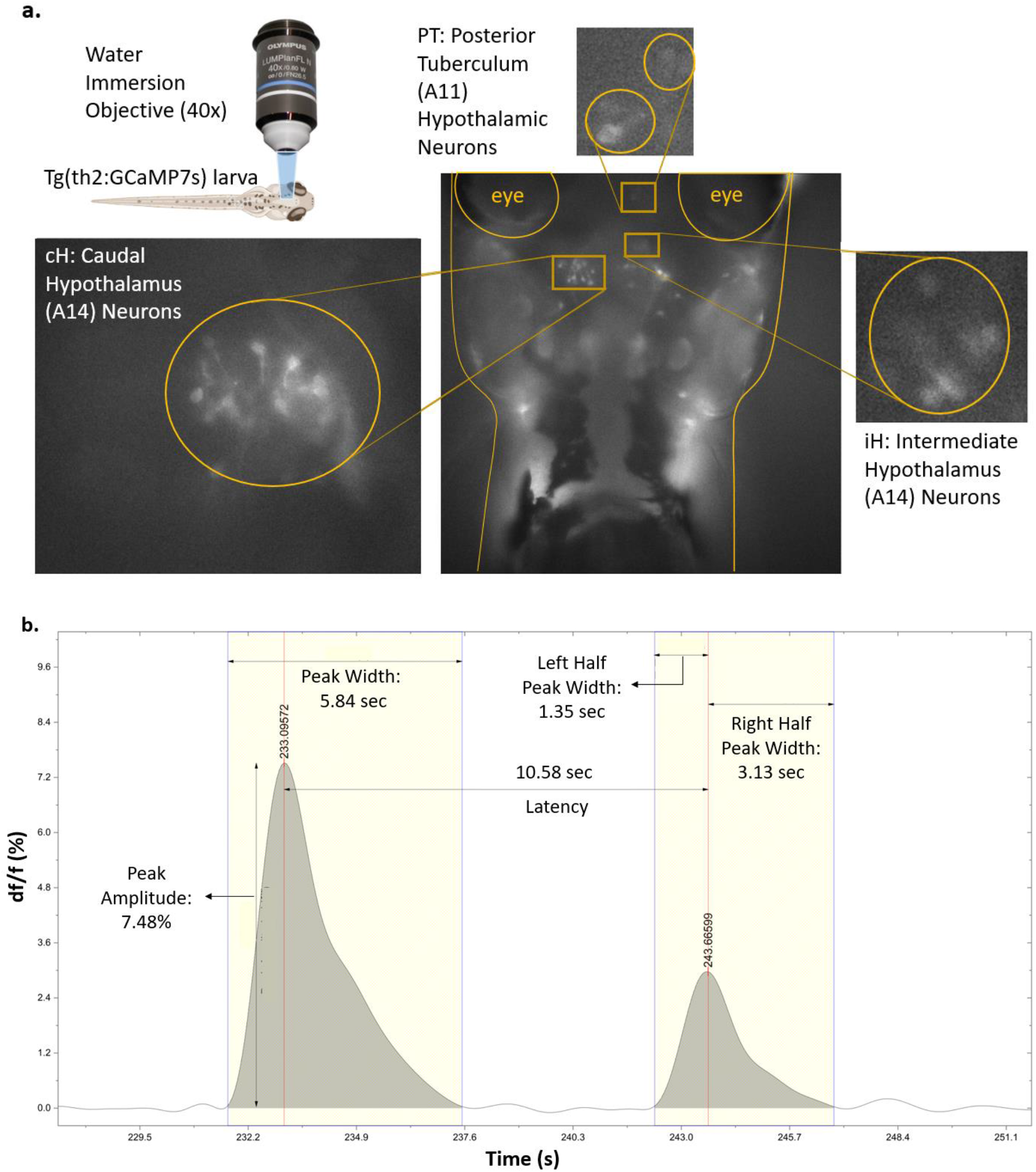
Larval imaging, dopamine neuron populations, and spike parameters. Calcium activity in three dopaminergic neuron populations (cH, iH, PT) in the hypothalamus of the transgenic larvae Tg(th2:GCaMP5) was analyzed with its characteristic parameters. **a.)** In the transgenic larvae, calcium activity in DA neurons in the caudal hypothalamic region, intermediate hypothalamic region, and the posterior tuberculum were recorded pre- and post-AMPH. The recording setup included a 40x water immersion objective connected to an epifluorescence microscope, and the individual larvae were paralyzed and immobilized in a drop of agarose. **b.)** In the calcium activity, five different parameters (df/f, peak width, left half peak width, right half peak width, and latency) were analyzed. The peak amplitude/normalized fluorescence change in the calcium activity was denoted by df/f; peak width denotes the activity duration of individual peaks, left half peak width denotes the peak rise time, and right half peak width represents the falling peak of the calcium peak. The latency between peaks denotes the time interval between two subsequent peaks.

### Post Processing

The data from ImageJ was extracted as intensity vs frames, and frames were converted into time in Microsoft Excel. Newly generated time vs intensity data were exported to OriginPro (Version 2022, OriginLab Corporation, Northampton, MA, USA). The Data were divided population-wise and were first smoothened (F) using an FFT filter (window points = 30). A baseline (Fo) was created from the smoothened intensity data using an inbuilt function called ‘peak analysis’ from the analysis tab using an asymmetric least squares smoothing algorithm by varying parameters suitable for the data. The baseline data was subtracted from smoothened intensity data and divided by baseline data (df/f F-Fo/Fo). The df/f values were opened again with the peak analysis function, and the peak integration function was used to obtain peak count and left and right half peak width values. The latency was calculated by subtracting two subsequent time points from the peaks (peak height threshold: ≥ 10% of df/f; local points = 2) (Fig. 1b).

### Statistical Analysis

All data were statistically analyzed using OriginPro. Data was checked for normality and was found to be rejecting normality. For Baseline analysis, a non-parametric test, the Mann-Whitney-U test for independent samples, was used between two data sets. Kruskal-Wallis ANOVA was performed for dose-dependent and latency analysis, followed by Dunn’s post-hoc analysis. Correlation analysis was performed using Spearman Correlation. All data are shown with significance factors (* *p* < 0.05, ** *p* < 0.01, *** *p* < 0.001, and **** *p* < 0.0001), and no significance was shown for non-significant p-value (>0.05). Box plot limits are represented as Q1-Q3:25%-75%, and whiskers are represented by outliers (1.5QRThe black horizontal line within the box represents the median with values mentioned on the right side of the line, and the black dot represents the mean of the data.

## Results

### Food deprivation increases baseline dopamine activity in the hypothalamus

Feeding states alter dopaminergic activity in animals. Here, we examined how food deprivation alters the dopaminergic activity in different brain regions in zebrafish larvae. In Fig. 2 a-c., we compared the peak amplitude between ad libitum (AL) and food-deprived (FD) groups in three distinct dopaminergic neuron populations in the hypothalamus (cH: caudal Hypothalamus; iH: intermediate Hypothalamus; PT: Posterior Tuberculum). With the statistical comparison of the peak amplitude data, we found that food deprivation increased the fluorescence activity in all three DA neuron populations with prominence in cH neurons (median ΔF/Fo: AL: 0.94%; FD: 2.17%; Mann-Whitney-U test: p<0.0001) followed by iH neurons (median ΔF/Fo: AL: 0.98%; FD: 1.46%; Mann-Whitney-U test: p<0.0001) and PT neurons (median ΔF/Fo: AL: 0.76%; FD: 1.21%; Mann-Whitney-U test: p<0.0001). Furthermore, we also investigated the peak width, which represents the time the DA neurons took to show one whole spike in the form of a calcium trace (Fig. 2 d-f). We did not find a significant width difference in peaks between AL and FD larvae in cH neuron (median peak width: AL: 4.41s; FD: 4.63s; Mann-Whitney-U test: p>0.05) however in both iH (median peak width: AL: 5.57s; FD: 6.24s; Mann-Whitney-U test: p<0.001) and PT neurons (median peak width: AL: 4.31s; FD: 4.55s; Mann-Whitney-U test: p<0.05), the peaks were significantly wider in FD state. In Fig. 2 g-i, we analyzed the left-half peak width of the calcium trace peaks that signify the time taken by the neuron activity peaks to reach the maximum amplitude beginning from the resting state. Here, statistical significance in left half peak width between AL and FD state can be found in all three DA neuron populations significantly in iH neurons (median peak width: AL: 0.714s; FD: 0.79s; Mann-Whitney-U test: p<0.0001), PT neurons (median peak width: AL: 0.766s; FD: 0.841s; Mann-Whitney-U test: p<0.0001). Similarly, we also compared the difference in right-half peak width, showing the time the calcium peaks took to reach the resting state from the maximum amplitude. The right half peak width was higher in the FD state in all three DA neurons. The difference was greatest in iH population (median peak width: AL: 0.82s; FD: 0.93s; Mann-Whitney-U test: p<0.0001) and then in PT neurons (median peak width: AL: 0.97s; FD: 1.06s; Mann-Whitney-U test: p<0.001) and lowest in cH neurons (median peak width: AL: 0.97s; FD: 1.05s; Mann-Whitney-U test: p<0.05).

**Figure 2.**
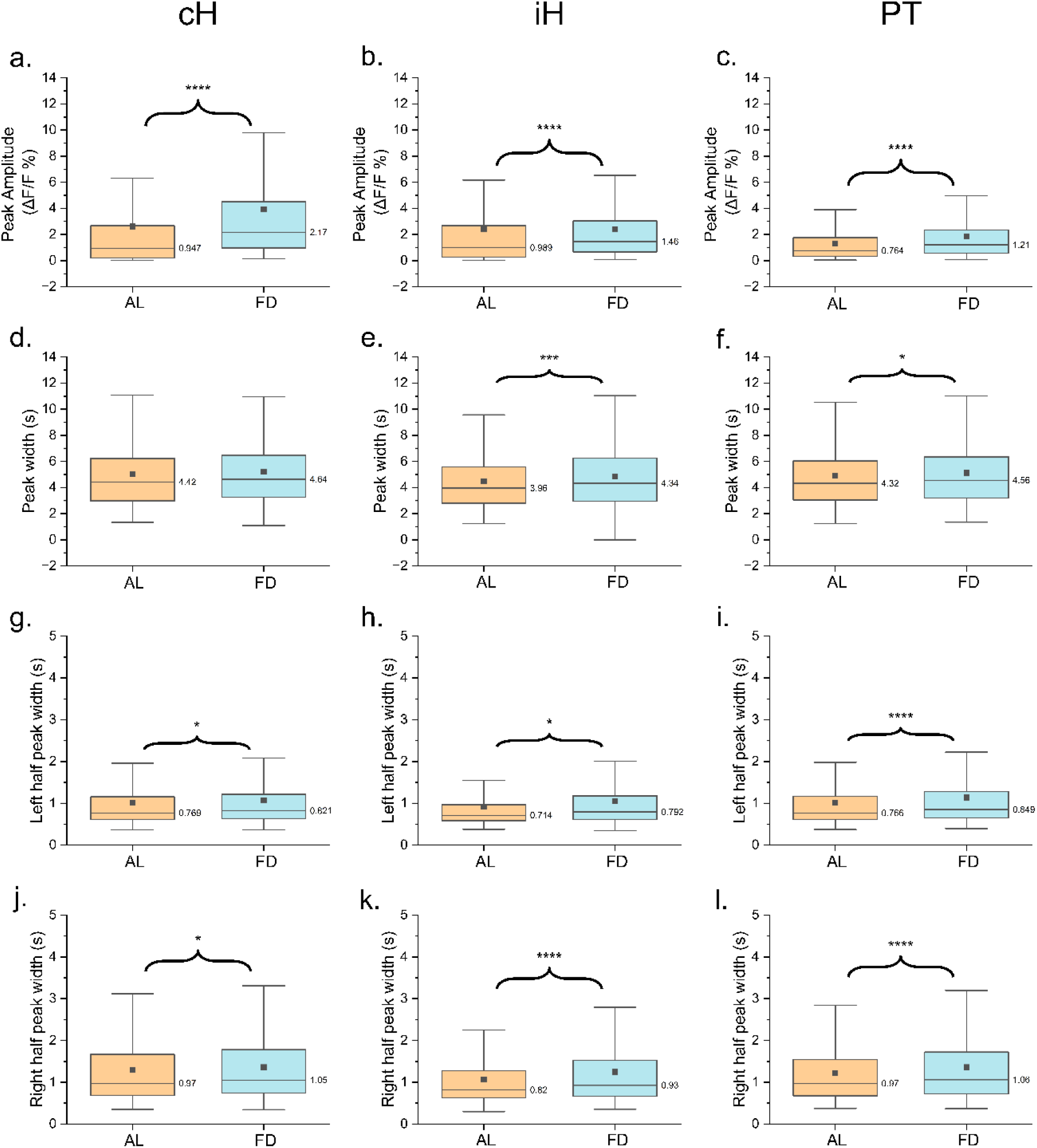
Baseline activity analysis between states. Baselines between Ad libitum (AL, n=20 fish) and Food-deprived (FD, n=18 fish) states were compared in four different calcium imaging data parameters that are peak amplitude (df/f)%, peak width, left half peak width, and right half peak width in three hypothalamic dopaminergic populations (caudal Hypothalamus (cH), intermediate Hypothalamus (iH) and Posterior Tuberculum (PT)) The data in all distribution is non-normal and thus tested using Mann-Whitney U test for independent data. **a-c)**. Fluorescence amplitude in neurons in untreated (baseline) state was significantly higher in the FD state than in the AL state in cH, iH, and PT (p<0.0001) **d-f)**. The fluorescence signal peak in baseline groups was significantly wider in FD larvae in iH (p<0.001) and PT neurons only (p<0.51) and mildly wider in cH and PT neurons. **g-i)**. Baseline left half peak widths were significantly wider in the FD state in all DA populations (cH: p<0.05); iH, PT: p<0.0001). **j-l)**. Right half peak width was significantly higher in FD than in AL in all DA populations (cH: p<0.05; iH: p<0.0001; PT: p<0.001). Box plot limits are represented as Q1-Q3:25%-75%, and whiskers are represented by outliers (1.5QR). The black horizontal line within the box represents the median with values mentioned on the right side of the line, and the black dot represents the mean of the data.

### Varying AMPH doses selectively exhibited increased calcium trace amplitude in DA neuron populations in caloric states

Amphetamine targets dopaminergic neurons and increases their activity in the brain to induce a rewarding effect. Here, we investigated the effects of two doses of amphetamine (0.7uM and 1.5uM) to gain insight into the reward-related functioning of the recently identified DA populations in the hypothalamus in zebrafish larvae. We compared the response in these neurons between AL and FD groups in the above-mentioned AMPH doses. We first compared the calcium trace peak amplitude (fluorescence change) between feeding states in two doses using Kruskal-Wallis ANOVA (Fig. 3 a-c). Comparison made in cH neurons (Fig. 3a) showed an overall significance [*χ*_*3*_^*2*^ = 113.44; n = 405 (AL_0.7), n = 387 (FD_0.7), n = 361 (AL_1.5), n = 349 (FD_1.5); p<0.0001]. Post-hoc comparison indicated that both doses in the FD state showed an increased neuron fluorescence relative to the AL state in cH population {median ΔF/Fo: 0.7uM (AL: 1.22%; FD: 1.56%), p<0.001; 1.5uM (AL: 0.819%; FD: 2.1%), p<0.0001}. In Fig. 3b, overall significant effect in iH neurons was observed [*χ*_*3*_^*2*^ = 129.60; n = 498 (AL_0.7), n = 485 (FD_0.7), n = 449 (AL_1.5), n = 424 (FD_1.5); p<0.0001]. Planned comparison showed a higher peak amplitude in FD state compared to AL state was observed only at 1.5uM dose {median ΔF/Fo: 0.7uM (AL: 1.12%; FD: 1.09%), p>0.05; 1.5uM (AL: 0.40%; FD: 1.3%), p<0.0001}. Similar to the other two populations, Kruskal-Wallis ANOVA conveyed an overall significant difference in PT neurons [*χ*_*3*_^*2*^ = 104.68; n = 522 (AL_0.7), n = 513 (FD_0.7), n = 491 (AL_1.5), n = 379 (FD_1.5); p<0.0001]. Here, the peak amplitude of the calcium traces was found to be higher in FD larvae than in AL larvae at 1.5uM dose only {median ΔF/Fo: 0.7uM (AL: 0.76%; FD: 0.91%), p>0.05; 1.5uM (AL: 0.43%; FD: 0.8%), p<0.0001} (Fig. 3c).

**Figure 3.**
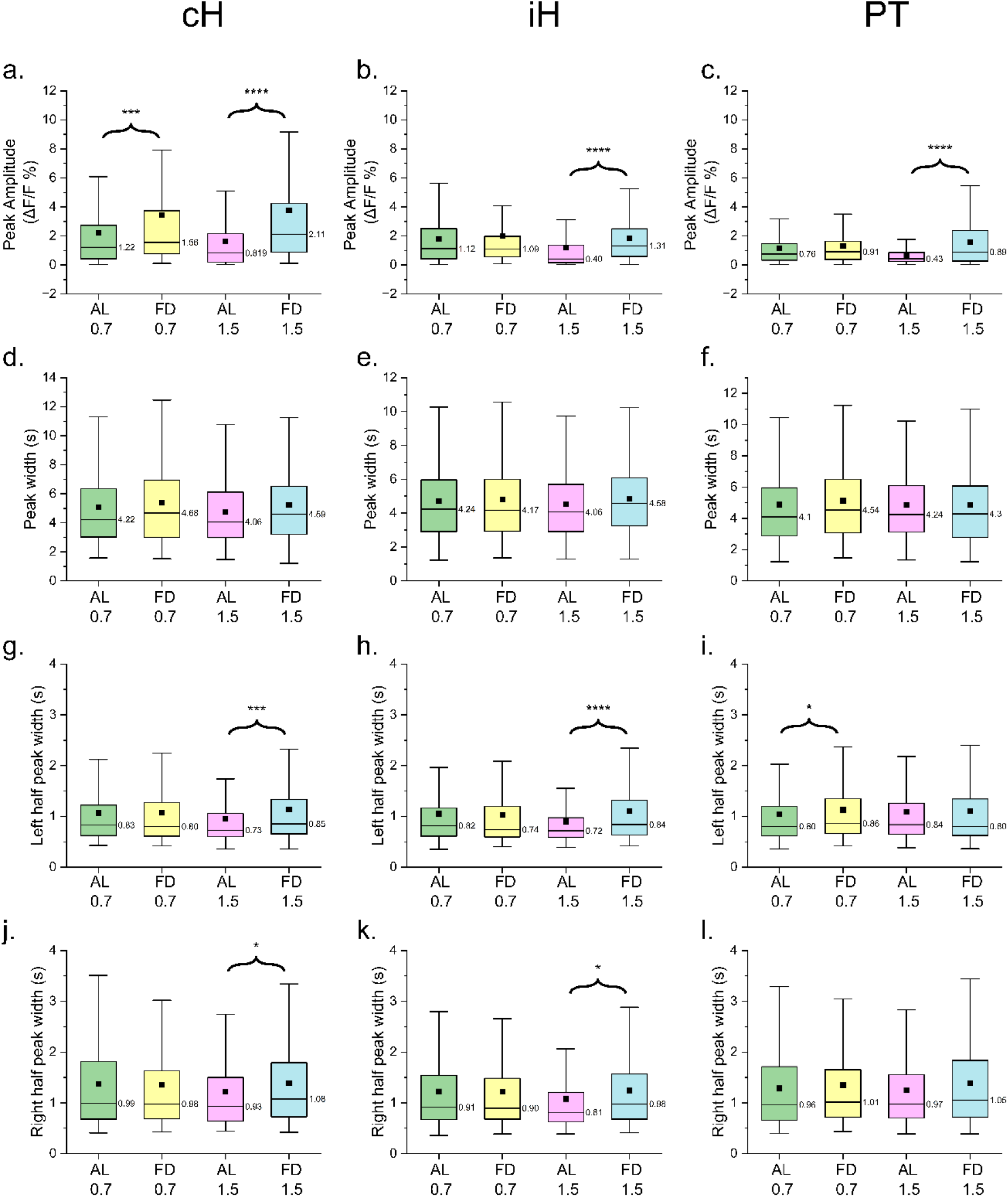
Dose-dependent analysis between states. All four parameters were statistically analyzed at two different doses of amphetamine [0.7uM (AL: n=11; FD: n=10 fish) and 1.5uM (AL: n=9; FD: n=8 fish)] and compared between AL and FD states using Kruskal-Wallis ANOVA for independent data followed by Dunn’s post-hoc comparison. Comparisons were made between AL and FD states in 0.7uM and 1.5uM doses of AMPH in all three DA neuron populations. **a-c)**. Both doses (0.7uM and 1.5uM) increased cH fluorescence activity (0.7uM: p<0.001; 1.5uM: p<0.0001) in cH neurons in FD state, and only 1.5uM AMPH dose increased neuron fluorescence in iH and PT populations (p<0.0001). **d-f)**. Overall peak width did not change significantly across doses and states in all three DA populations. **g-i)**. Left half peak width was significantly larger in FD states only at 1.5uM AMPH dose in cH (p<0.001) and iH (p<0.0001) and in AL state in PT neurons at 0.7uM dose (p<0.05). **j-l)**. The right half peak was wider in the FD state at 1.5uM in cH and iH population only (p<0.01). Box plot limits are represented as Q1-Q3:25%-75%, and whiskers are represented by outliers (1.5QR). The black horizontal line within the box represents the median with values mentioned on the right side of the line, and the black dot represents the mean of the data.

### Food deprivation potentiated hypothalamic DA activity by increasing the rising and falling duration of calcium peaks post-AMPH treatment

We investigated whether varying the AMPH dose affects the duration of neuron activity by measuring the whole peak width, left half peak width (peak rising fluorescence amplitude), and right half peak width (peak declining fluorescence amplitude) in individual calcium traces. In Fig. 3 d-e, we statistically analyzed the difference in whole peak width (resting-peak-resting amplitude). We did not find statistical significance in the peak width in any of the three DA neuron populations cH [*χ*_*3*_^*2*^= 7.70; n = 404 (AL_0.7), n = 386 (FD_0.7), n = 361 (AL_1.5), n = 350 (FD_1.5); p>0.05] (Fig. 3d), iH [*χ*_*3*_^*2*^= 6.15; n = 498 (AL_0.7), n = 485 (FD_0.7), n = 449 (AL_1.5), n = 424 (FD_1.5); p>0.05] (Fig. 3e) and in PT [*χ*_*3*_^*2*^ = 5.50; n = 521 (AL_0.7), n = 498 (FD_0.7), n = 491 (AL_1.5), n = 379 (FD_1.5); p>0.05] (Fig. 3f). Post-hoc comparison did not show a significant change between AL and FD states in cH {median peak width: 0.7uM (AL: 6.35s; FD: 6.92s), p>0.05; 1.5uM (AL: 6.13s; FD: 6.51s), p>0.05}, iH {median peak width: 0.7uM (AL: 5.96s; FD: 5.98s), p>0.05; 1.5uM (AL: 5.69s; FD: 6.08s), p>0.05} and PT {median peak width: 0.7uM (AL: 4.09s; FD: 4.53s), p>0.05; 1.5uM (AL: 4.24s; FD: 4.29s), p>0.05}. All three DA neuron populations exhibited a mild but non-significant increase in peak amplitude in FD state post AMPH treatment in all doses.

Upon analyzing left half peak width in Fig. 3g-i, which represents the time duration between resting and highest amplitude of the calcium peak, overall significant change was observed using Kruskal-Wallis ANOVA [*χ*_*3*_^*2*^ = 16.61; n = 404 (AL_0.7), n = 386 (FD_0.7), n = 361 (AL_1.5), n = 350 (FD_1.5); p<0.001] (Fig. 3g) and posthoc comparison also showed significance between AL and FD groups at 1.5uM dose only in PT {median peak width: 0.7uM (AL: 0.82s; FD: 0.79s), p>0.05 1.5uM (AL: 0.72s; FD: 0.85s), p<0.001}. Similarly, significance was also found in iH neurons between two caloric states [*χ*_*3*_^*2*^ = 24.91; n = 498 (AL_0.7), n = 485 (FD_0.7), n = 449 (AL_1.5), n = 424 (FD_1.5); p<0.0001] with planned comparison indicating difference between physiological states at 1.5uM dose {median peak width: 0.7uM (AL: 0.81s; FD: 0.73s), p>0.05; 1.5uM (AL: 0.71s; FD: 0.83s), p<0.0001} (Fig. 3h). In PT neurons, although overall significant difference was observed [*χ*_*3*_^*2*^ = 8.28; n = 522 (AL_0.7), n = 498 (FD_0.7), n = 491 (AL_1.5), n = 379 (FD_1.5); p<0.001], contrarily, post-hoc test showed the difference between caloric states only at 0.7uM dose {median peak width: 0.7uM (AL: 0.80s; FD: 0.86s), p<0.05; 1.5uM (AL: 0.83s; FD: 0.80s), p>0.05} (Fig. 3i). In cH and iH neuron populations, significantly higher left half peak width was observed in food-deprived groups post 1.5uM AMPH dose whereas same trend was shown by PT neurons but at 0.7uM dose depicting the neuron activity to take significantly longer to reach the maximum amplitude.

The right-half peak width was analyzed to measure the time the calcium trace peak took to decline from the highest to its resting state. In Fig. 3j, Kruskal-Wallis ANOVA test expressed the dosage effect in cH neurons [*χ*_*3*_^*2*^ = 9.14; n = 404 (AL_0.7), n = 386 (FD_0.7), n = 361 (AL_1.5), n = 350 (FD_1.5); p<0.05] and post-hoc comparison revealed a significant difference between states at 1.5uM dosage {median peak width: 0.7uM (AL: 0.98s; FD: 0.97s), p>0.05; 1.5uM (AL: 0.92s; FD: 1.07s), p<0.05}. Dose effect in iH neurons was also observed to be significant in Fig. 3k [*χ*_*3*_^*2*^ = 18.41; n = 498 (AL_0.7), n = 485 (FD_0.7), n = 449 (AL_1.5), n = 424 (FD_1.5); p<0.001]. FD larvae exhibited a significant increase at 1.5uM dosage when pairwise comparisons were done {median peak width: 0.7uM (AL: 0.91s; FD: 0.89s), p>0.05; 1.5uM (AL: 0.81s; FD: 0.97s), p<0.001}. Contrarily, significance was neither induced by dosage in PT neurons for right-hand peak width neurons [*χ*_*3*_^*2*^ = 4.74; n = 522 (AL_0.7), n = 498 (FD_0.7), n = 491 (AL_1.5), n = 378 (FD_1.5); p>0.05] nor by posthoc comparison {median peak width: 0.7uM (AL: 0.96s; FD: 1.01s), p>0.05; 1.5uM (AL: 0.97s; FD: 1.05s), p>0.05} (Fig. 3l). Right-hand peak width data shows that the FD state increased the time taken by the neuron activity to drift down from highest to resting amplitude causing the neuron activity to stay longer.

### Feeding states impaired the neuron correlation between DA neuron pairs independently and associatively with AMPH doses

In Fig 4, we measured the dopamine neuron activity correlation between neuron pairs at baseline level (without AMPH) and two different doses. The correlation conveys how strongly the neuron activity is correlated between different DA neuron pairs and how feeding states and AMPH affect this correlation. Although the correlation coefficients calculated here are weak when their individual values were considered, we only made a relative comparison in this analysis. Firstly, we compared the correlation between neuron pairs in both states at baseline level (Fig. 4 a-b). In the AL state (Fig. 4a), We observed a significantly high correlation between all pairs. A relatively strong positive correlation was observed between cH-PT neuron activity (Spearman’s R Correlation: 0.1, p<0.01), which was followed by the cH-iH pair (Spearman’s R Correlation: 0.086, p<0.05) however, a negative correlation was shown by iH-PT neuron pair (Spearman’s R Correlation: -0.089, p<0.05). Interestingly, when correlation was measured between neuron pairs in FD state (Fig. 4b), although correlation coefficients were significant in between all pairs, we found a relatively stronger positive correlation between the iH-PT pair (Spearman’s R Correlation: 0.25, p<0.0001) which was contrary to the observation made in AL state. A relatively weak positive correlation was observed between the iH-cH pair (Spearman’s R Correlation: 0.21, p<0.0001), followed by a weaker but significant positive correlation between the cH-PT pair (Spearman’s R Correlation: 0.15, p<0.001)

**Figure 4.**
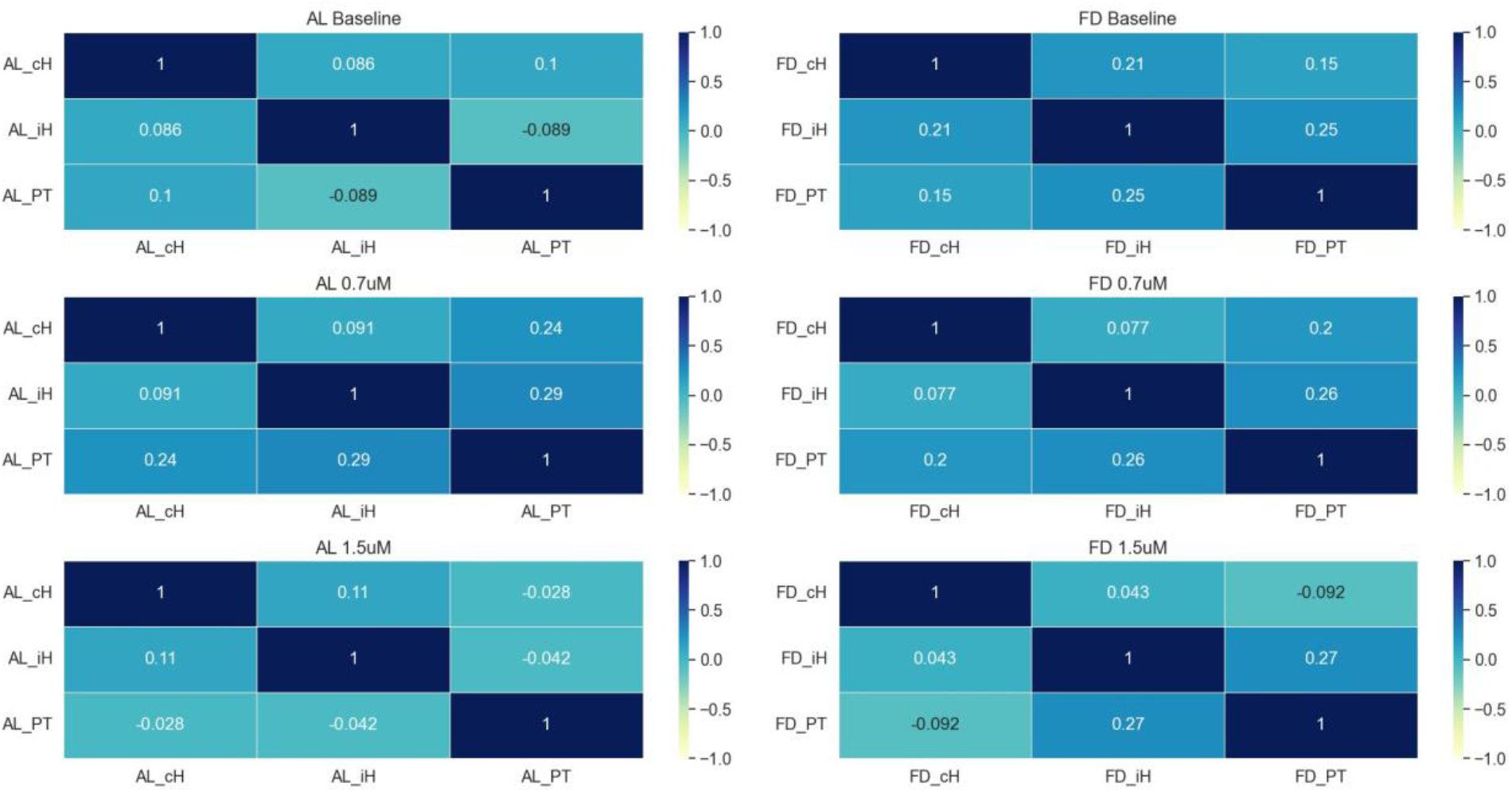
Calcium peak amplitude (df/f) correlation heatmap. Correlation analysis of DA neuron activity among all three DA populations in AL and FD states was performed in baseline and two AMPH doses. **a-b). Baseline:** Neuron activity correlation between cH and PT was highest in AL larvae, followed by cH and iH neurons, whereas PT-iH neurons showed a negative correlation. FD larvae showed the highest neuron activity correlation between iH-PT neurons, followed by cH-iH neurons, and the least correlated pair was cH-PT. **c-d). 0.7uM:** In the AL state, highly correlated pairs at lower AMPH doses were PT-iH and PT-cH, showing a noticeable increase at this dosage. cH-iH pairs were the least correlated here. Correlation in neuron activity in FD larvae showed an identical trend. **d-e). 1.5uM:** At a higher AMPH dose of 1.5uM, cH-iH activity correlation in AL larvae was the highest among all DA populations. At this dosage, cH-PT and iH-PT neuron pairs showed a negative correlation in the activity. In the FD state, neuron activity was correlated between the iH-PT neuron pair followed by the cH-iH pair. Similar to the AL state, the FD state induced a negative correlation between the cH-PT pair.

When neuron activity correlation was measured in larvae after AMPH treatment, a significant change and reduction in correlation was observed in some pairs. In Fig 4c, when AL larvae were treated with 0.7uM AMPH, iH neurons were strongly correlated with PT neurons (Spearman’s R Correlation: 0.29, p<0.0001). The positive correlation between cH and PT neurons was relatively weaker (Spearman’s R Correlation: 0.24, p<0.0001), and the least non-significant positive correlation was expressed by cH-iH pair (Spearman’s R Correlation: 0.091, p>0.05). Larvae in the FD state treated with 0.7uM AMPH (Fig. 4d) expressed a strong positive correlation between the iH-PT neuron pair (Spearman’s R Correlation: 0.26, p<0.0001). A significantly strong positive correlation was also observed in (Spearman’s R Correlation: 0.20, p<0.0001), and the least positive and non-significant correlation was exhibited by the cH-iH pair (Spearman’s R Correlation: 0.077, p>0.05). At a higher AMPH dose (1.5uM), neuron activity correlation between all neuron pairs in the AL state significantly reduced (Fig. 4e). Strongest and significant positive correlation was seen between cH and iH (Spearman’s R Correlation: 0.11, p<0.01). Activity correlation was found to be reduced and statistically non-significant between the cH-PT pair (Spearman’s R Correlation: -0.028, p>0.05) and iH-PT neuron pair (Spearman’s R Correlation: -0.042, p>0.05). Contrary to the AL state, FD state post AMPH administration indicated the strongest positive correlation in this dosage in the iH-PT neuron pair (Spearman’s R Correlation: 0.27, p<0.0001), and the least positively correlated pair was iH-cH (Spearman’s R Correlation: 0.043, p>0.05) and was non-significant. Between cH and PT neurons, similar to the AL state at 1.5uM dose, activity correlation was negative and significant (Spearman’s R Correlation: -0.092, p<0.01).

### Feeding affected the calcium spike latency dose-dependently in selective DA populations

Latency in neuronal calcium spikes depicts the time elapsed between two subsequent adjacent peaks and possesses an inverse relationship with neuron activity frequency. We looked into neuron calcium trace latency and compared it pre- and post-AMPH at both doses and all three DA neuron populations. Interestingly, we found the outcomes contrary to what we saw in peak amplitude and width investigation dose-dependently. At 0.7uM dose, we did see an overall significant change in latency after AMPH administration in cH neurons after Kruskal-Wallis ANOVA [*χ*_*3*_^*2*^ = 16.11; n = 391 (AL Baseline), n = 394 (AL_0.7), n = 367 (FD Baseline), n = 376 (FD_0.7); p<0.01] (Fig. 5a). No significant latency change post-AMPH was observed except a mild increase in AL state and decrease in FD state {median difference: AL Baseline-AL_0.7: 0.33s, p>0.05; FD Baseline-FD_0.7: -0.11s, p>0.05}. However, at 1.5uM dose (Fig. 5b), along with an overall significance with ANOVA [*χ*_*3*_^*2*^= 10.75; n = 302 (AL Baseline), n = 352 (AL_1.5), n = 348 (FD Baseline), n = 342 (FD_1.5); p<0.05] planned comparison also showed a significant decrease in latency in cH neurons in AL state but a mild non-significant increase was observed in FD state {median difference: AL Baseline-AL_1.5: -0.68s, p<0.05; FD Baseline-FD_1.5: 0.15s, p>0.05}. In Fig. 5c, iH neurons at 0.7uM neither showed an overall significant effect of state [*χ*_*3*_^*2*^ = 5.40; n = 445 (AL Baseline), n = 488 (AL_0.7), n = 448 (FD Baseline), n = 475 (FD_0.7); p>0.05] nor latency change after AMPH treatment in both AL {median difference: AL Baseline-AL_0.7: -0.13s, p>0.05; FD Baseline-FD_0.7: -0.41s, p>0.05}. Latency in iH neurons at 1.5uM dose (Fig. 5d) followed the same trend as it did at 0.7uM dose. Statistical significance was not observed either with ANOVA [*χ*_*3*_^*2*^ = 5.86; n = 412 (AL Baseline), n = 440 (AL_1.5), n = 405 (FD Baseline), n = 416 (FD_1.5); p>0.05] and post-hoc test in both caloric states {median difference: AL Baseline-AL_1.5: -0.22s, p<0.05; FD Baseline-FD_1.5: 0.19s, p>0.05}. PT neuron at 0.7uM dose did not express an overall significant change in latency [*χ*_*3*_^*2*^ = 3.75; n = 490 (AL Baseline), n = 512 (AL_0.7), n = 498 (FD Baseline), n = 488 (FD_0.7); p>0.05] (Fig. 5e). Pairwise comparison also failed to show significance after AMPH administration {median difference: AL Baseline-AL_0.7: -0.2s, p>0.05; FD Baseline-FD_0.7: 0.29s, p>0.05}. PT neurons expressed a significant overall effect at 1.5uM [*χ*_*3*_^*2*^ = 26.73; n = 407 (AL Baseline), n = 482 (AL_1.5), n = 343 (FD Baseline), n = 371 (FD_0.7); p>0.05]. Pairwise comparison indicated a significant decrease in latency in the AL state but not in the FD state {median difference: AL Baseline-AL_1.5: -0.92s, p<0.0001; FD Baseline-FD_1.5: -0.45s, p>0.05}. The results showed a decrease in latency (increased neuron activity) when larvae were fully fed (AL state), indicating that food reward interferes with the rewarding properties of AMPH to induce a dose-dependent change in DA neuron activity in these hypothalamic populations.

**Figure 5.**
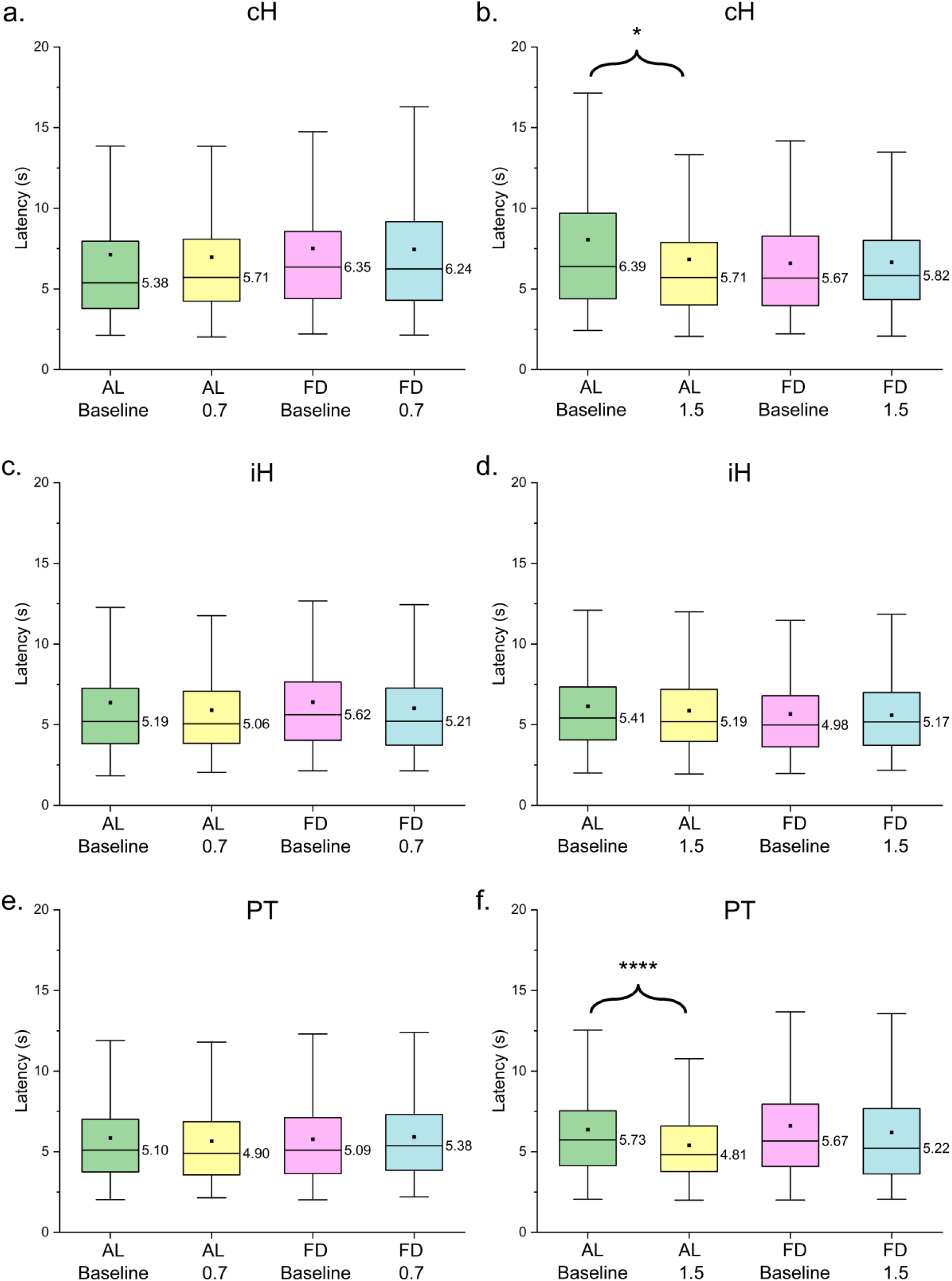
Inter-spike interval (Latency) comparison. Latency was statistically analyzed at two different doses of amphetamine [Baseline and 0.7uM (AL: n=11; FD: n=10 fish) and Baseline and 1.5uM (AL: n=9; FD: n=8 fish)] between paired subjects. Paired comparison between baselines and doses in AL and FD states using Kruskal-Wallis ANOVA for dependent data followed by Dunn’s posthoc comparison. Pairwise comparisons were made between baselines and their respective doses in AL and FD states (pre- and post-AMPH) in all three DA neuron populations. **a-b). cH:** Latency significantly decreased in cH neurons at 1.5uM AMPH dose in FD state only (Median difference: 0.15; p<0.05). **c-d). iH:** No significant difference pre- and post-AMPH was observed in latency in iH neurons in both doses. **e-f). PT:** Decreased latency was only observed in the AL state at 1.5uM dose (Median difference: 0.92; p<0.0001) and remained non-significant everywhere else. Box plot limits are represented as Q1-Q3:25%-75%, and whiskers are represented by outliers (1.5QR). The black horizontal line within the box represents the median with values mentioned on the right side of the line, and the black dot represents the mean of the data.

## Discussion

Using zebrafish larvae, this study investigated A11 (PT) and A14 (cH, iH) type dopamine (DA) neurons in the hypothalamus (Barrios et al., 2020; Haehnel-Taguchi et al., 2018). Generally, DA activity is influenced by the rewarding nature of food and drugs in the mesolimbic pathway, which initiates from the ventral tegmental area (VTA) and is projected into the nucleus accumbens (NAc), a part of the striatum in the mammalian brain (Holly & Miczek, 2016). In zebrafish, research shows the innervations of DA neurons from the posterior tuberculum (PT) into the subpallium brain region in zebrafish, which is a teleost equivalent of mammalian striatum (Rink & Wullimann, 2002). However, no evidence exists to identify the mammalian equivalent of the complete limbic reward pathway in the zebrafish brain. Moreover, how the interference of food and drug rewards affects the dopaminergic pathway has yet to be well understood. Thus, there is a considerable knowledge gap regarding the identification of reward circuitry in this animal model and the effects of complexing the drug reward with different feeding regimes on the known DA neuron populations. Therefore, to fill this gap, in this novel study, we measured and evaluated the individual and interactive effects of food deprivation and different doses of amphetamine (AMPH) on three distinct dopaminergic neuron populations in the hypothalamus of zebrafish larvae and compared the responses with ad libitum fed larvae.

Zebrafish larvae are born with a yolk sac, which includes a supply of essential nutrition that serves as food only till 4 pdf and requires external feeding upon reaching 5 pdf (Sant & Timme-Laragy, 2018). We used 6dpf larvae with an 18-hour acute overnight food-deprivation period to obtain dopaminergic activity recordings. According to our findings, food deprivation alone increased neuron activity in all three DA populations. Our results agree with the outcomes in previous studies that were performed in acutely fasted rodents and acute central ghrelin injection (Anderberg et al., 2016; Roseberry, 2015). Dopaminergic neurons receive synapses from various neuron populations in the brain (Watabe-uchida et al., 2012). One such neuron population is Agouti-related protein/Neuropeptide-Y (AgRP/NPY) neurons in the hypothalamus, which get stimulated when ghrelin (a gut hormone that is secreted during starvation) binds to its receptors on AgRP/NPY neuron surface (Essner et al., 2017; Khelifa et al., 2021). Food deprivation increases AgRP/NPY mRNA expression (Bi et al., 2003). In addition, the direct incremental effect of ghrelin on dopaminergic neurons has been observed via its binding onto ghrelin receptors (Abizaid, 2009; Jerlhag & Egecioglu, 2010; Quarta et al., 2009). These results altogether indicate that A11 and A14 DA populations are affected by hunger and probably receive synaptic inputs directly or indirectly from AgRP/NPY neurons and are also affected by ghrelin release. Thus, the presence of AgRP/NPY innervations in and ghrelin receptors on zebrafish A11 and A14 neurons can be investigated in the future.

We also reported that neuron activity amplitude was relatively higher in the FD state than in the AL state with both doses and was comparatively greater with the higher dose in all three DA populations. Food deprivation and AMPH-mediated increase in DA neuron activity could have arisen from decreased insulin levels in food-deprived states. Studies show that dexamethasone (an insulin blocker) injection and food deprivation independently increase NPY mRNA expression, and FD-mediated NPY release increases DA activity (Rezitis et al., 2022; Sato et al., 2005). In mice, amphetamine treatment significantly decreases fasting blood glucose, and reduced blood glucose further lowers insulin levels (Fruehwald-schultes et al., 2000; Y. Zhang et al., 2018). Although studies in the context of insulin-hunger-AMPH are scarce, especially in zebrafish, more than 70% of glucose and insulin regulatory genes studied in zebrafish and humans are conserved and show a similar regulation pathway (Y. Zhang, Qin, et al., 2018). Thus, the reduction in insulin caused by food deprivation and its further decrease by AMPH treatment might have led to an increase in NPY levels, which probably caused a surge in DA activity in FD larvae. Investigating the connection among glucose, insulin, food deprivation, and amphetamine can shed more light on their collective impact on DA activity in mammals and zebrafish. The aforementioned study showing the innervation of DA neurons from PT into the subpallium and the increase in dopaminergic neuron activity in PT along with cH and iH in food-deprived states points to PT being the primary reward-related region. From our findings and these previous observations, it would be reasonable to assume that the differences in brain structure may have functionally compensated for the rewarding properties by providing a part of the central rewarding function to the PT-striatum circuitry in this teleost. However, this assumption holds unless the reward-related DA populations are identified. Despite the limited study of cH and iH DA neuronal projections, an increase in their activity also points to their yet unknown innervations into the zebrafish striatum besides PT DA neurons and could be explored in the future to understand their function in eliciting hedonic reward.

After peak amplitude, we measured the activity duration of calcium peaks (peak width) followed by peak rise time (left half peak width) and peak fall time (right half peak width). We found that baseline peak width, rise, and fall time were higher in the FD state in all three neuron populations, especially with significance in iH and PT neurons. Since we have used calcium imaging, the peak duration (peak width) corresponds to the calcium ion channel opening response that facilitates the flux of calcium ions in the neurons. These channels open in response to the binding of neurotransmitters to the neuron’s NMDA and AMPA receptors (Pchitskaya et al., 2018). These two receptors harbor the binding site for glutamate, and its binding on these receptors activates them and opens calcium ion channels (Traynelis et al., 2010). A study shows that acute starvation leads to increased glutamate concentration in the brain (Mourek et al., n.d.). This starvation-mediated glutamate increase may have led to the activation of ion channels for an extended period compared to the fed state, leading to wider calcium signaling peaks in the neurons. Further, activation of AMPA and NMDA receptors increases dopamine neuron firing (Zakharov et al., 2016). NMDA receptors in this previous study were shown to affect DA neuron activity more than AMPA receptors, whereas activation of both increased DA activity more than their individual activation. Thus, the comparatively low activation rate of both the receptors or absence of NMDA receptors in baseline activity cH DA neurons in our study could be the reason for the less significant increase in peak width in the FD state. At baselines, although the calcium peak activity duration was longer in the FD state, there could have been a delay in calcium ion flux that can be observed in the time taken by the peak to rise to its highest (Left half) and again to the lowest (right half) amplitude. Further studies are needed to investigate the role of starvation in ion flux and channel dynamics in DA neurons.

Afterward, treatment with amphetamine at a lower dose decreased the difference in peak width between AL and FD states. A significant change was observed in peak rise and fall time at a higher AMPH dose only in A14 neurons. This change could be occurring because AMPH increases NMDA receptor-mediated synaptic currents in dopamine neurons, whereas it decreases AMPA-mediated synaptic activity in the same (Li et al., 2017). Moreover, glycine is another amino acid that acts as a co-agonist with glutamate in NMDA receptor activation, and its concentration also increases during starvation (Becker & Betz, 2012; Felig et al., 1969). AMPH alters glycine levels dose-dependently (Mora, 1993). Thus, glycine levels may have been significantly lowered at the doses used here and failed to evoke a significant change in peak width between both states. Additionally, Glycine-Glutamate-AMPH interaction could have impaired the calcium ion movement compared to baseline. Various homologs of both receptors have been identified in zebrafish (Horzmann & Freeman, 2016). However, the arena of food deprivation-AMPH interaction with these receptors still needs to be explored in zebrafish and mammals. A detailed investigation is required to understand the functionality of these homologous receptors to study the changes occurring in A11 and A14 DA activity in feeding states and with stimulant treatment.

In the AL and FD states, we performed df/f correlation analysis between DA neuron pairs pre- and post-AMPH treatment. The baseline calcium activity (df/f) correlation between all DA pairs, both within the A14 neurons and between A14 and A11 neurons, was relatively higher in the food-deprived state than in the ad libitum state. A11 neurons receive inputs from several nuclei, including the parabrachial nucleus (PBN), which is important in modulating feeding-related homeostasis (Koblinger et al., 2014; Z. Zhao et al., 2023). In zebrafish, the Secondary Gustatory-General Viscerosensory Nucleus (SGN/V) is present as the mammalian equivalent of PBN containing food/taste sensory neurons and projects into the regions of the posterior tuberculum (Henriques et al., 2019; Yáñez et al., 2022). Here, calretinin (CR) is expressed predominantly in SGN, which is a part of the periventricular hypothalamus in PT and caudal/intermediate hypothalamic regions in zebrafish brains, and food deprivation increases its expression (Hua et al., 2018; Mueller, 2012). Studies show that the presence of CR in striatal DA neurons in different animal species exhibits a variety of effects of CR in DA neurons and its precursor L-DOPA (Isaacs et al., 1997; Mura et al., 2000; Petryszyn et al., 2016). Yet, the effects of CR on DA excitability remained largely unexplored. Dopaminergic activity is also affected by AgRP neurons, and in the zebrafish brain, AgRP neurons are located in the periventricular hypothalamus while projecting throughout the hypothalamic regions (Shainer et al., 2017). Additionally, hypothalamic AgRP neurons are relatively closer to A14 neurons. Therefore, an increase in DA activity simultaneously via SGN-CR in A11 and AgRP in both A11 and A14 DA neurons could be a possible reason for the relatively higher correlation in activity between both DA types in food deprivation. Calcitonin gene-related peptide (CGRP) is a gut protein that is present in both mammals and zebrafish, and its expressive neurons are present in PBN and A11 neurons in mammals (Kuil et al., 2021; Milet et al., 2018; Z. Zhao et al., 2023). CGRP has been shown to suppress food intake in animals in fed states (Sanford et al., 2019). This suggests that the fed state might have induced the release of CGRP, and it also increases DA neuron activity either directly from A11 neurons or via a neuron pathway other than calretinin (Rahimi et al., 2017). Contrarily, a recent study showed that ablation of AgRP neurons led to the impairment of dopaminergic neuron activity in rodents, and AgRP neuron activity generally decreases in the fed state (Reichenbach et al., 2022). This contradictory activity in dopaminergic neurons in the A11 and A14 regions may have led to a decrease in correlation relative to what we have observed in the food-deprived state.

More interestingly, with AMPH treatment, the activity correlation increased in the AL state relative to the FD state at both doses, and the increment was relatively more noticeable at the higher dose. This altered correlation from amphetamine treatment could stem from its effect on a peptide called protein kinase C (PKC) in PBN that mediates taste transduction (Varkevisser & Kinnamon, 2023). Research shows that both food and amphetamine increase PKC activity, and an independent study shows a PKC-mediated increase in dopamine activity (Inoguchi et al., 1939; Krivanek, 1997; Mutanen et al., 2000). However, food and AMPH both have been shown to inhibit AgRP neuron activity, and AgRP ablation decreased dopamine neuron excitation in rodents (Alhadeff et al., 2019; Reichenbach et al., 2022). These two findings are contradictory, but identical modulation in PKC and AgRP-mediated conditions express an increased correlation in the AL state. The function of PKC has been identified in zebrafish only recently for its involvement in fat and glucose metabolism (Sun et al., 2021). Hence, there is a need to perform further studies about its participation in modulating DA neuron activity along with the use of stimulants. Moreover, the difference in activity correlation within and between neuron populations also showcases the ability of feeding states to leverage different pathways in modulating the neuron activity correlation. This can be observed especially when activity between iH and PT in the FD state exhibited the highest correlation; it went to its lowest negative value in the AL baseline state and in both AL and FD states at higher AMPH doses.

Upon investigating calcium peak latency (time duration between two adjacent peaks), the lower dose of AMPH did not induce any significant change in peak latency. Whereas at higher dose, the AL state was significantly affected and a decreased latency can be observed in cH and PT neurons. These outcomes exhibit an increase in the frequency of calcium peak occurrence. In this pre-and post-AMPH treatment analysis at both doses, we found that latency was significantly lowered at a higher dose in the AL state only in cH and PT neurons. We also observed impairment in latency/spike rate outcomes, which is still being debated as randomness in the neuron activity occurring naturally, which is by design and depends on the type of stimulus given (Fiorillo et al., 2014). Yet, the change in calcium dynamics arising from feeding states and AMPH interaction could be explained since calcium ions play an important role in evoking action potentials in the neurons (Gleichmann & Mattson, 2011). Intuitively, to observe decreased latency in neuron activity, calcium ion influx, and outflux must increase in frequency so that the spikes can occur more frequently. Along with other cells, neurons contain Adenosine Triphosphate (ATP), consumed by the cells to generate energy (Zampese et al., 2022). These ATP molecules are utilized more in the fed state than in the fasted state and remove excess cytosolic calcium ions from the cells, whereas the fasted state retains ATP reserves (R. Zhao & Zhao, 2021). Moreover, amphetamine decreases intracellular ATP by increasing its utilization (Brown & Yamamoto, 2003). Thus, reduced latency (increased spike rate) in the AL state could arise from increased utilization of ATP, leading to a higher calcium ion removal rate from the neurons. Further, the faster influx of calcium ions may have occurred due to the acute calcium ion gradient formed from its quick removal. Additionally, a study also showed ATPs’ role in increased striatal dopaminergic activity which could explain an increased calcium peak amplitude in food-deprived states due to higher ATP reserves (Zlhang et al., 1995). Recently, zebrafish also altered ATP metabolism towards diet (Roy et al., 2021). Studying energy metabolism in the future could help us comprehend the role of feeding states-AMPH-ATP consumption and could increase our understanding of changes in DA activity related to energy homeostasis.

## Conclusion

Our results show that the dopaminergic neurons exhibited increased activity in all three hypothalamic populations when subjected to acute fasting alone and with AMPH treatment relative to the fed state. Exploring this cohesive effect of caloric state-AMPH on DA neurons further in the future will help devise interventional strategies for controlling the DA activity in individual populations by varying diet and stimulant doses. The caloric state-AMPH treatment also altered the correlation between populations. This could also be studied to understand how different DA populations communicate when subjected to rewarding stimuli. Our results regarding peak rising and falling time, which also increased with hunger-AMPH interaction, could be a topic of future study, which could shed light on the mechanism of receptor, channel, and ion dynamics related to DA neurons and could be used to modulate the DA activity.

## Acknowledgments

We thank Dr. Adam Douglass for providing the Tg(th2:GCaMP7s) transgenic line. We also thank John Jutoy for helping with fish room maintenance.

## Author Contributions

Conceptualization: E.E.J.; Methodology: E.E.J. and P.B.; Investigation: E.E.J. and P.B; Formal Analysis: P.B.; Visualization: P.B.; Writing: E.E.J., and P.B; Original Draft: P.B.; Review and Editing: E.E.J.; Funding Acquisition: E.E.J; Resources: E.E.J.; Supervision: E.E.J.

## Data Availability Statement

The data will be made available from the corresponding author upon reasonable request.

